# Bioactive constituents of *Verbena officinalis* alleviate inflammation and enhance killing efficiency of natural killer cells

**DOI:** 10.1101/2021.10.22.465498

**Authors:** Xiangdong Dai, Xiangda Zhou, Rui Shao, Renping Zhao, Archana K. Yanamandra, Zhimei Xing, Mingyu Ding, Junhong Wang, Han Zhang, Yi Wang, Qi Zheng, Peng Zhang, Bin Qu, Yu Wang

## Abstract

Natural killer (NK) cells play a key role in eliminating pathogen-infected cells. *Verbena officinalis* (*V. officinalis*) has been used as a medical plant in traditional and modern medicine, exhibiting anti-tumor and anti-inflammation activities, but its roles in immune responses still remains largely elusive. In this work, investigated the regulation of inflammation and NK functions *by V. officinalis* extract (VO-extract). In an influenza virus infection mouse model, oral administration of VO-extract alleviated lung injury, promoted maturation and activation of NK cells residing in the lung, and decreased the levels of inflammatory cytokines (IL-6, TNF-α and IL-1β) in the serum. We further analyzed the impact of five bioactive components of VO-extract on NK killing functions. Among them, Verbenalin enhanced NK killing efficiency significantly as determined by real-time killing assays based on plate-reader or high-throughput live-cell imaging in 3D using primary human NK cells. Further investigation showed that treatment of Verbenalin accelerated killing processes by reducing the contact time of NK cells with their target cells without affecting NK proliferation, expression of cytotoxic proteins, or lytic granule degranulation. Together, our findings reveal that low doses of *V. officinalis* can achieve a satisfactory anti-inflammation effect against viral infection *in vivo*, and *V. officinalis* regulates activation, maturation and killing functions of NK cells. NK killing efficiency is enhanced by Verbenalin from *V. officinalis*, suggesting a promising potential of verbenalin to fight viral infection.

## 1. Introduction

*Verbena officinalis* L. (*V. officinalis*), also known as common vervain, is a medicinal herb, widespread throughout the globe, mainly in the temperate climate zone^[1]^. In China, *V. officinalis* is widely distributed in the southern part of the Yellow River and has been widely used not only as traditional Chinese medicine for the treatment of rheumatism, bronchitis, depression, insomnia, anxiety, liver and gallbladder diseases ^**Error! Reference source not found.**^, but also in food and cosmetics with a long-standing record for validated safety^**Error! Reference source not found.**^. Flavonoids, terpenoids, phenolic acids, phenylpropanoids and iridoids are its mainly identified bioactive constituents^[2]^. Recent reports suggest that *V. officinalis* has various scientifically proven activities, such as anti-oxidation, anti-bacteria, anti-fungi, and anti-inflammation properties^[6]^.

Inflammation is a immune response triggered by numerous factors, including virus, bacteria, and transformed cells. During inflammation, permeability of blood vessels is enhanced, facilitating recruitment of immune cells to the inflammation site. Recruited immune cells release cytokines to further activate and recruit other effector immune cells. Inflammatory responses play an essential role in fighting pathogens. However, uncontrolled inflammatory responses can lead to severe consequences for example organ functional failure especially in lung and life-threatening cytokine storm syndrome. Innate immune cells are the main players to initiate inflammation responses.

In the innate immune system, natural killer (NK) cells are specialized immune killer cells, which play a key role in eliminating tumorigenic and pathogen-infected cells. After viral infection, NK cells are recruited to the lungs and play an essential role in the immune response to fight pathogens. Several studies highlight the pivotal role of NK cells in the control of infection of influenza virus H1N1. Defects in NK cell activity or depletion of NK cells result in delayed viral clearance and increased morbidity and mortality ^**Error! Reference source not found.**^. However, there are also examples in which NK cells exacerbate morbidity and pathology during lethal dose influenza virus infection in mice ^**Error! Reference source not found.**^, indicating that overactivation of NK cells may lead to undesirable effects. In addition, NK cells play important roles in bridging the innate and adaptive immune responses to viral infection ^**Error! Reference source not found.**^.

In this work, we investigated the anti-inflammation effect of VO-extract *in vivo* with low (0.5 g/kg) and high (1 g/kg) doses using a viral infection mouse model. We found that both doses significantly reduced viral infection-induced release of inflammatory cytokines (TNFα, IL-6, and IL-1β). Of note, the low dose exhibited a better protection of the lung tissue and could induce higher level of NK activation. Further analysis of bioactive constituents from *V. officinalis* revealed that Verbenalin substantially enhanced killing efficiency of NK cells.

## 2. Materials and Methods

### 2.1. *Preparation of* VO-extract

The *Verbena officinalis L*. was obtained from Anhui Zehua China Pharmaceutical Slices Co., Ltd. The whole plants of *Verbena officinalis L*. (4.5 kg) were extracted with 4.5 L 70% ethanol for 2 hours by refluxing extraction repeated for three times. The combined extract was filtrated with ceramic membranes. The filtration was then concentrated with by vacuum evaporation apparatus at a temperature not exceeding 45 °C. Then the extract was lyophilized to obtain the VO-extract powder.

### 2.2. UPLC-Q-TOF-MS analysis

An Agilent 1290 UHPLC system (Agilent Technologies Inc., Palo Alto, CA, USA) coupled to an Agilent 6520 Q-TOF instrument with electrospray ionization (ESI) source was used for the quantitative analysis. The ACQUITY UPLC^®^ BEH C18(2.1×150 mm, 1.7μm; Waters, Milford, MA, USA) was used for chromatographic separation. The mobile phase consisted of 0.1% aqueous formic acid (A) and methanol (B). The elution conditions involved holding the starting mobile phase at 95% A and 5% B and applying a gradient of 5% A and 95% B for 35 min. The flow rate was 0.3 mL/min, and the injection volume for all the sample was 2 μL. Experiments were performed in positive and negative ESI mode with the following parameters:temperature of ESI, 100 °C; collision energy, 10 V; collision pressure, 135V; fragmentor voltage, 135 V; nebulizer gas, 40.0 psi; dry gas, 11.0 L/min at a temperature of 350 °C; scan range, m/z 100-1700.

### 2.3. Mice and virus

6–10 week old female C57BL/6 mice(20 to 25 g body weight) were purchased from Beijing Vital River Laboratory Animal Technology Co., Ltd. (Beijing, China), and housed in standard microisolator cages in a centralized animal care facility. Animal care and experimental procedures were performed in accordance with experimental-animal guidelines. Mice were given ad libitum access to food and water, and subject to 12 h light/dark cycling. All mice were adapted to the environment for seven days before the study. Virus (H1N1) were frozen at −80°C.

### 2.4. Virus infection

Mice were anesthetized with isoflurane and intranasally inoculated with H1N1 in 40 μL PBS. We inoculated mice with 100 PFU H1N1. The number of mice in each group ranged from 6 to 8. The mice in the Vehicle group were intranasally challenged with 40 μL PBS. All the animal experiments were approved and made to minimize suffering and to reduce the number of animals used.

### 2.5. VO-extract administration

VO-extract (VL: 492mg/kg; VH: 984mg/kg) was administered by oral gavage once daily, 3 days for the duration of treatment. VO-extract was freshly prepared each day and stored at 4 °C. The mice in the Vehicle group were orally administered with distilled water simultaneously. On the third day, the mice were sacrificed, and their blood and lung were collected for further analysis. Meanwhile, mice were monitored and body weight changes were recorded.

### 2.6. Flow cytometry assay

After blocking the receptor with anti-CD16/CD32, extracellular markers were stained, cells were fixed. These cells were incubated with specific surface-binding antibodies for 30 min at 4°C. Samples were analyzed using BD FACScalibur and FlowJo software. Cells were gated according to forward scatter and side scatter, and cell types were identified by phenotype as follows: for NK cells:CD45, NK1.1, CD11b, CD69.

### 2.7. Hematoxylin–eosin (HE) staining

For the assay of lung pathological changes, the mice were sacrificed on the third day, and the lung tissues of the infected mice were collected, fixed in 10% buffered formalin, and embedded in paraffin. Each tissue was cut into 4 μm sections and stained with hematoxylin and eosin. Subsequently, Lung injury was evaluated according to a quantitative scoring system assessing infiltration of immune cells, thickening of alveolar walls, disrupted lung parenchyma^[14]^. Scoring was performed by three researchers independently according to standard protocols.

### 2.8. Detection of IL-6, TNF-α and IL-1β levels in serum

Blood samples were centrifuged at 3000 rpm and 4 °C for 15 min to obtain the sera. The contents of IL-6, TNF-α and IL-1β in the sera were measured by ELISA following the manufacturer’s instructions and by using a microplate reader. The contents of these cytokines were determined by establishing standard curves.

### 2.9. NK Cell preparation and cell culture

Primary human NK cells were isolated from peripheral blood mononuclear cells (PBMCs) of healthy donors using Human NK Cell Isolation Kit (Miltenyi). The isolated NK cells were cultured in AIM V medium (ThermoFischer Scientific) with 10% FCS and 100 U/ml of recombinant human IL-2 (Miltenyi). K562 and K562-pCasper cells were cultured in RPMI-1640 medium (ThermoFischer Scientific) with 10% FCS. For K562-pCasper cells, 1.25mg/ml G418 was added. All cells were kept at 37 °C with 5% CO_2_.

### 2.10. Real-time killing assay

Real-time killing assay was conducted as reported previously^**Error! Reference source not found.**^. Briefly, target cells (K562 cells) were loaded with Calcein-AM (500 nM, ThermoFisher Scientific) and settled into a 96-well plate (2.5×10^4^ target cells per well). NK cells were subsequently added with an effector to target (E:T) ratio of 2.5:1 if not otherwise mentioned. Fluorescence intensity was determined by GENios Pro micro-plate reader (TECAN) using the bottom-reading mode at 37°C every 10 min for 4 hours. Target lysis (t) % = 100 × (Flive(t)-Fexp(t))/(Flive(t)-Flysed(t)). (F: fluorescence intensity)

### 2.11. 3D killing assay and live cell imaging

Briefly, target cells (K562-pCasper cells) were embedded into 2 mg/ml of pre-chilled neutralized Bovine type I collagen solution (Advanced Biomatrix) in a 96 well plate. The collagen was solidified at 37°C with 5% CO_2_ for 40 min. NK cells were subsequently put on top of collagen as effector cells. The cells were visualized using ImageXpress Micro XLS Widefield High-Content Analysis System (Molecular Devices) at 37 °C with 5% CO_2_. For 3D killing assay, as described previously^[18]^, the killing events were visualized every 20 min for 36 hours, and live target cell numbers were normalized to hour 0 based on area. For live cell imaging to determine time required for killing and the average kills per NK cell, the cells were visualized every 70 sec for 14 hours and tracked manually. Image J software was used to process and analyze the images.

### 2.12. Proliferation assay

To examine proliferation, freshly isolated primary human NK cells were labelled with CFSE (1 μM, ThermoFischer Scientific) and then stimulated with recombinant human IL-2 in presence of Verbenalin at indicated concentrations for 3 days. Fluorescence was determined with a FACSVerse™ flow cytometer (BD Biosciences) and analyzed with FlowJo v10 (FLOWJO, LLC).

### 2.13. Determination of cytotoxic protein expression

To test perforin and granzyme B expression, NK cells were fixed in pre-chilled 4% paraformaldehyde. Permeabilization was carried out using 0.1% saponin in PBS containing 0.5% BSA and 5% FCS. FACSVerse™ flow cytometer (BD Biosciences) was used to acquire data. FlowJo v10 (FLOWJO, LLC) was used for analysis.

### 2.14. CD107a degranulation assay

For degranulation assay, K562 cells were settled with vehicle-treated or Verbenalin-treated NK cells in the presence of Brilliant Violet 421™ anti-human CD107a (LAMP1) antibody (Biolegend) and GolgiStop™ (BD Biosciences). The incubation were carried out at 37°C with 5% CO_2_ for 4 hours. The cells were then stained with PerCP anti-human CD16 antibody (Biolegend) and APC mouse anti-human CD56 antibody (BD Biosciences) to define NK cells. Data was obtained with a FACSVerse™ flow cytometer (BD Biosciences) and was analyzed with FlowJo v10 (FLOWJO, LLC).

### 2.15. Statistical analysis

Data were analyzed using the SPSS version 19.0 and the GraphPad Prism software 5.0 and presented as mean ± standard deviation (SD). Significant differences among the multiple group comparisons were performed using one-way analysis of variance (ANOVA), and the ANOVA comparisons were analyzed through Tukey’s honest significant difference test, whereas data with a partial distribution were examined using the nonparametric Kruskal–Wallis test with Dunn’s multiple comparison as post-test. A p value < 0.05 indicated significant difference.

## 3. Results

### 3.1. VO-extract attenuates the acute lung damage induced by viral infection

Lung damage induced by infection of virus, for example influenza or SARS-CoV (severe acute respiratory syndrome coronavirus)-1/2 can cause severe breathing problems or even respiratory failure ^[19]^. Infection-induced acute lung damage can result in persist lung abnormalities such as pulmonary fibrosis, leading to long-term impairments in the respiratory system for the patients recovered from the infection ^[20]^. Infection-induced acute lung damages are, to a large extent, owed to a massive inflammation initiated by the overactivated immune system. The well-established anti-inflammation activity of *V. officinalis* prompted us to examine its effect on infection-induced lung damage. To this end, we infected C57BL/6J mice with an influenza virus A/PR8/34 (H1N1) and intragastrically administrated VO-extract upon viral infection once per day for 3 days (Figure 1A). We chose the low dose (0.5 g/kg) and the high dose (1 g/kg) based on the doses in previous studies in rats^[21]^. Body weight was monitored daily. Loss of weight was observed in the virus infected group, and administration of VO-extract significantly alleviated this infection-induced weight loss for both the low and the high dose (Figure 1B). Using H&E staining in lung sections, we next examined changes in alveolar morphology and immune cell infiltration on day 3 post viral infection. We observed massive infiltration of immune cells along with thickening of alveolar walls and disrupted lung parenchyma in virus-infected mice (Figure 1C, Virus group vs. Control group). These infection-induced symptoms in lung was considerably reduced by the administration of VO-extract for low and high doses (Figure 1C, Low/High group vs. Virus group). Intriguingly, the low dose seemed to achieve a better attenuation of the symptoms relative to the high dose (Figure 1C, Low group vs High group). Along the same line, viral infection-triggered release of the inflammatory cytokines (TNF-α, IL-1β and IL-6) was abolished by the administration of VO-extract to a comparable level as Control group (Figure 1D). No difference was identified between high and low doses (Figure 1D). Taken together, our findings indicate that VO-extract is a potent agent to dampen infection-induced acute lung damage and inflammatory response.

**Figure 1.**
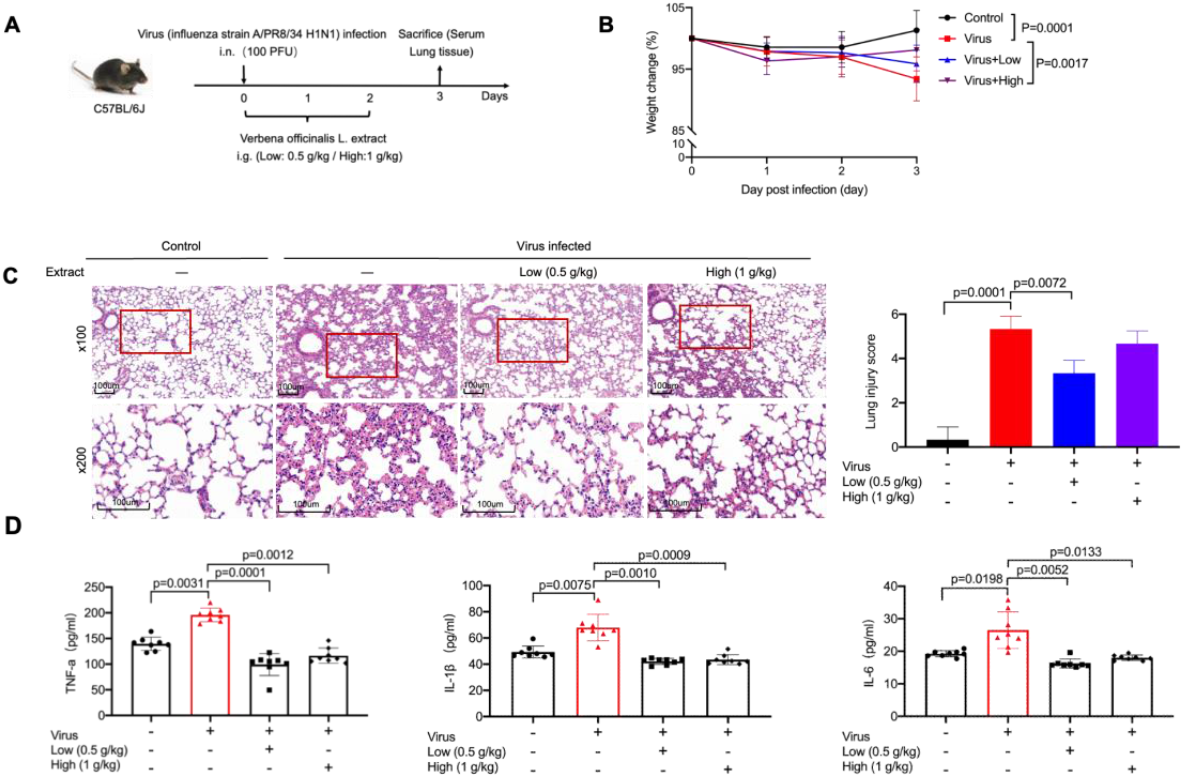
Analysis of the effects of VO-extract on infection-induced acute lung damage and inflammation *in vivo*. (A) C57BL/6J mice were challenged intranasally with influenza virus A/PR8/34 (H1N1) (100 PFU) on day 0, followed by oral administration with a single dose of VO-extract (low dose/Low: 0.5 g/kg; high dose/High: 1 g/kg) every day for three days. Mice were sacrificed on day 3. (B) Viral infection caused loss of body weight was ameliorated by VO-extract. Body weight of mice was measured every day for three days. n = 8 for each group. (C) Administration of VO-extract alleviated virus-induced inflammation in lung. Histological analysis of lung tissue was carried out on day 3. Two magnifications are shown: 100× (scale bars: 100 μm) and 200× (scale bars: 100 μm). One representation sample from each group is shown (n = 3). a total of 50 alveoli were counted on each slide at ×200 magnification (n=3). (D) Administration of VO-extract abolished viral infection triggered release of proinflammatory cytokines. Blood samples were taken on day 3. Cytokine concentration was determined using ELISA. SPSS version 19.0 and the GraphPad Prism software 5.0 were used for statistical analysis. Results are shown as mean±SD.

### 3.2. VO-extract promotes infection-induced maturation and activation of NK cells in the lungs

NK cells are key players to eliminate pathogen-infected cells. We next analyzed the impact of VO-extract on NK functions. For this purpose, we quantified the frequency and activation of the NK cells isolated from the lung tissue on day 3 post-infection. We found no alteration in the frequency of lung-residing NK cells by administration of VO-extract (Figure 2A, B). To further evaluate NK cell activation, we used surface markers CD11b and CD69, as expression of CD11b is positively associated with effector functions of murine NK cells^[23]^ and expression of CD69 is induced in activated NK cells and contribute to their cytotoxic functions^[24]^. We found that viral infection substantially enhanced the frequency of the CD11b^+^ NK subset, and this tendency was further elevated by administration of the low dose of VO-extract but not altered by the high dose (Figure 2A, C). A similar effect was also observed for the CD69^+^ NK subset (Figure 2A, D). These results indicate that only low dose of VO-extract promote viral infection-induced activation of NK cells.

**Figure 2.**
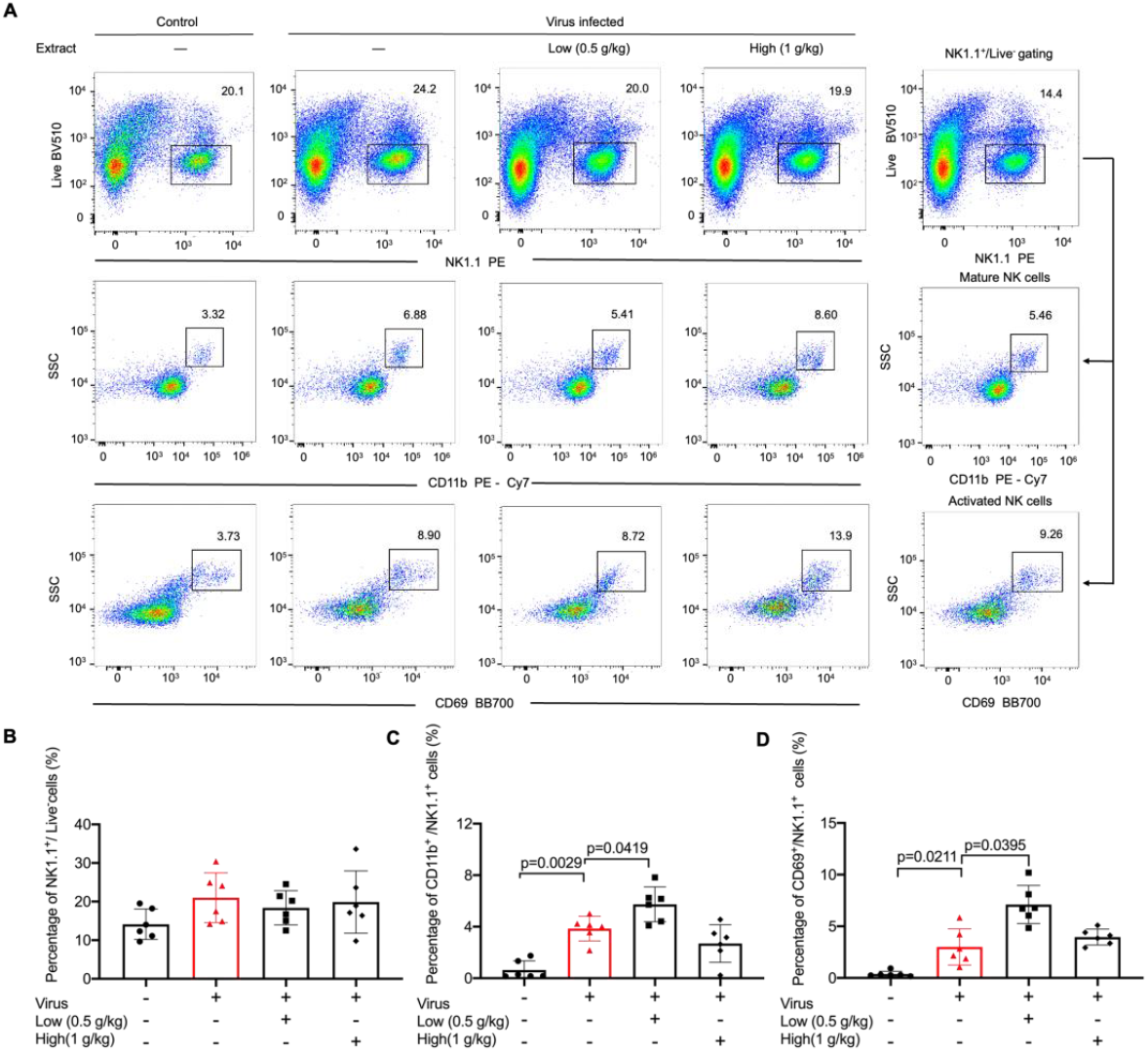
Low dose of VO-extract enhances NK activation in response to viral infection. C57BL/6J mice were challenged intranasally with virus (influenza virus A/PR8/34 (H1N1), 100 PFU) on day 0, followed by oral administration with a single dose of VO-extract (Low: 0.5 g/kg; High: 1 g/kg) every day for three days. Lung samples were collected on day 3. NK cells from lungs were examined using flow cytometry. SPSS version 19.0 and the GraphPad Prism software 5.0 were used for statistical analysis.

### 3.3. Identification of chemical composition from VO-extract by UPLC-Q-TOF-MS

To identify the active ingredients in VO-extract, we performed ultra-high performance liquid chromatography-quadrupole time-of-flight mass spectrometry (UPLC-Q-TOF-MS). A representative base peak chromatogram (BPC) of VO-extract based on the positive and negative ion modes is shown in Figure 3. A total number of thirteen ingredients were successful identified: 3,4-dihydroverbenalin, Verbeofflin I, Hastatoside, Verbenaloside, Quercetin, Acteoside, Luteolin, Isorhamnetin, Luteolin 7-O-β-gentiobioside, Isoacteoside, Leucosceptoside A, Apigenin, Kaempferol (Supplementary Table 1).

**Figure 3.**
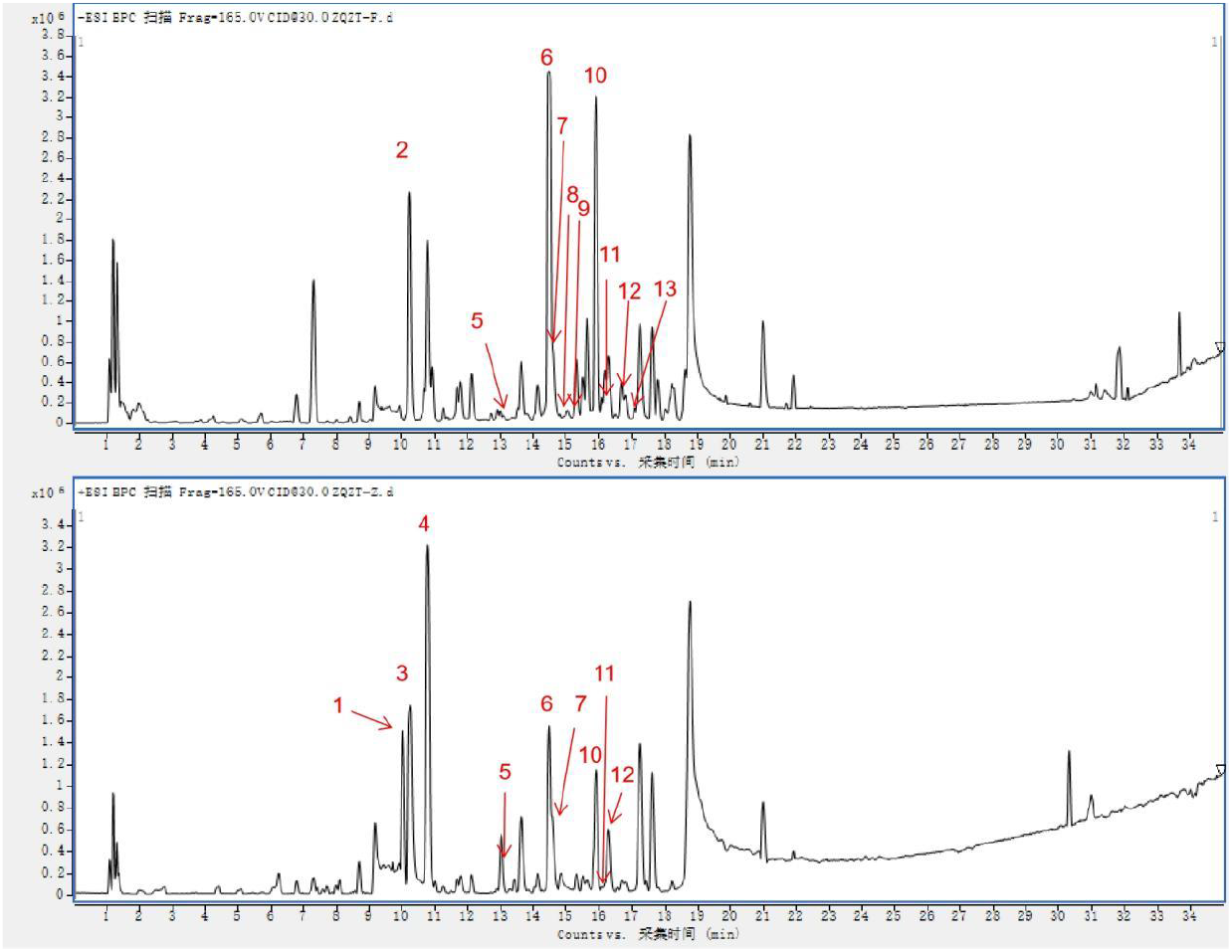
The chemical base peak intensity (BPI) chromatogram of key compounds characterization of VO-extract in positive ion mode and negative ion mode determined by UPLC-Q-TOF/MS. Identification No.: 3,4-dihydroverbenalin (1); Verbeofflin I (2); Hastatoside (3); Verbenaloside (4); Quercetin (5); Acteoside (6); Luteolin (7); Isorhamnetin (8); Luteolin 7-*O-β*-gentiobioside (9); Isoacteoside (10); Leucosceptoside A (11); Apigenin (12); Kaempferol (13).

### 3.4. Bioactive components of V. officinalis enhance NK killing efficiency

In order to verify the impact of bioactive constituents of *V. officinalis* on NK effector functions, we cultured primary human NK cells with the corresponding compound (10 μM and 30 μM) for three days in presence with IL-2. First, we analyzed killing kinetics of NK cells using a plate-reader based real-time killing assay ^**Error! Reference source not found.**^.

We found that Acteoside, Apigenin, and Kaempferol slightly reduced NK killing efficiency, whereas Verbenalin and Hastatoside enhanced NK killing efficiency (Figure 4A). Verbenalin exhibited the highest potency for elevation of NK killing efficiency (Figure 4A). We further verified the impact of Verbenalin on NK killing efficiency in 3D. Target cells expressing FRET-based apoptosis reporter pCasper were embedded in collagen matrix and NK cells were added from the top. Target cells were yellow when alive and turned green when undergoing apoptosis ^**Error! Reference source not found.**^. Results show that Verbenalin-treated NK cells exhibited significantly faster killing kinetics compared to their counterparts treated with vehicle. Thus, we conclude that among the bioactive constituents of *V. officinalis*, Verbenalin is able to increase NK killing efficiency under a physiologically relevant condition.

**Figure 4.**
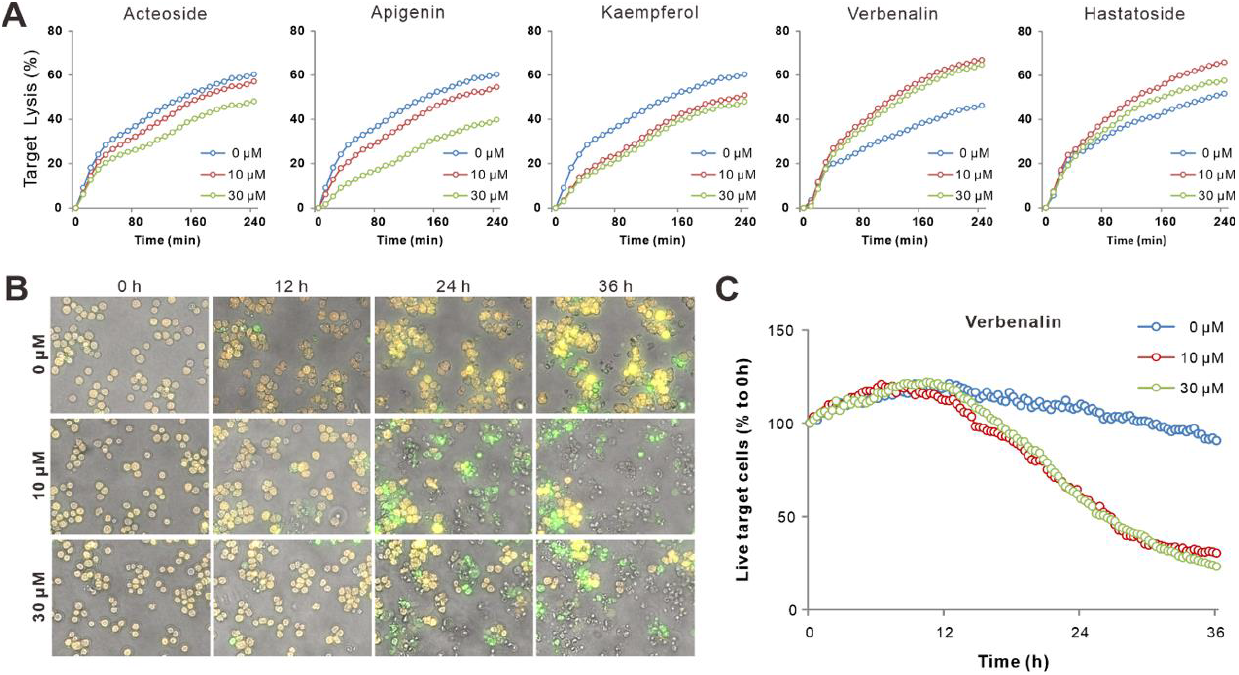
Bioactive constituents of Verbena differentially regulated NK killing efficiency. (A) Kinetics of NK killing affected by bioactive constituents of *V. officinalis*. were determined with real-time killing assay. (B, C) Verbenalin accelerates NK killing kinetics in 3D. Selected time points are shown in B and the change in live target cells is shown in C. Scale bars are 40 μm. One representative donor out of four is shown.

### 3.5. Verbenalin accelerated NK killing processes

Next, we sought for the underlying mechanisms of increase in NK killing efficiency by Verbenalin. We examined proliferation, expression of cytotoxic proteins (perforin and granzyme B), and degranulation of lytic granules. None of those processes were affected by Verbenalin (Figure 5A-C). We then analyzed the times required for killing by visualization of the killing events with high-content imaging setup every 70 sec for 12 hours (Figure 6A, Movie 1). The time between initiation of NK/target contact and target cell apoptosis was analyzed for randomly chose NK cells. The quantification shows that the time required for killing was considerably reduced by Verbenalin-treatment (Figure 6B). Concomitantly, on average, the numbers of target cells killed per NK cell was almost doubled for Verbenalin-treated NK cells relative to their vehicle-treated counterpart (Figure 6C). Notably, it is reported that a portion of NK cells can serve as serious killers, which are able to kill several target cells in a row ^**Error! Reference source not found.**^. We thus also analyzed the frequency of serial killers (one NK killed more than one target cell), single killers (one NK killed only one target cell) and non-killers (NK cells did not kill any target cells). We found that Verbenalin-treatment substantially enhanced the portion of serial killers and concomitantly reduced the portion of non-killers (Figure 6D). Taken together, our results suggest that Verbenalin potentiates NK cell activation upon target recognition, shortening the time required to initiate destruction of target cells, therefore enhancing killing efficiency of NK cells.

**Figure 5.**
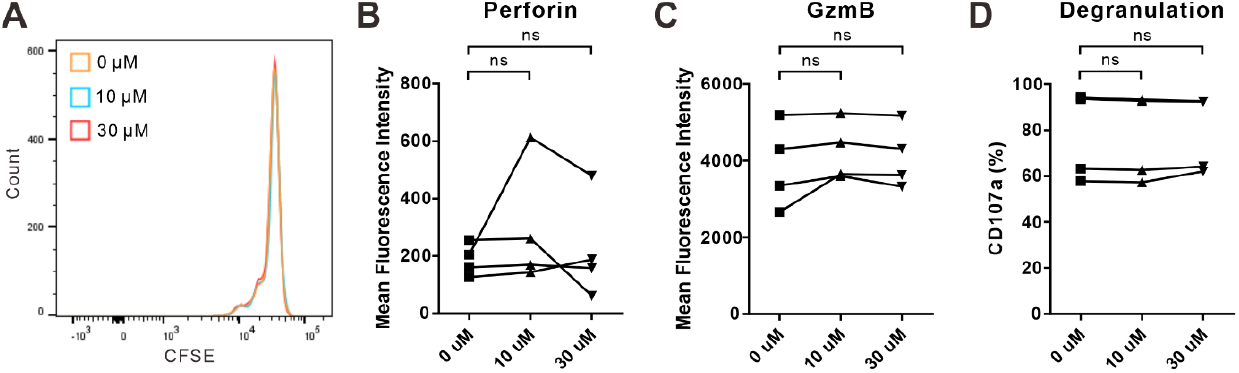
Proliferation and lytic granule pathway of NK cells were not affected by Verbenalin. Primary human NK cells were stimulated with IL-2 in presence of Verbenalin with indicated concentrations for 3 days prior to experiments. (A) Proliferation of NK cells. One representative donor out of four is shown. (B, C) Expression of cytotoxic proteins. Expression of perforin (B) and granzyme B (C) was determined by flow cytometry. Results are from four donors. (D) Release of lytic granules was determined with CD107a degranulation assay. Results are from four donors. ns: not significant. Paired Student’s t-test was used for statistical analysis.

**Figure 6.**
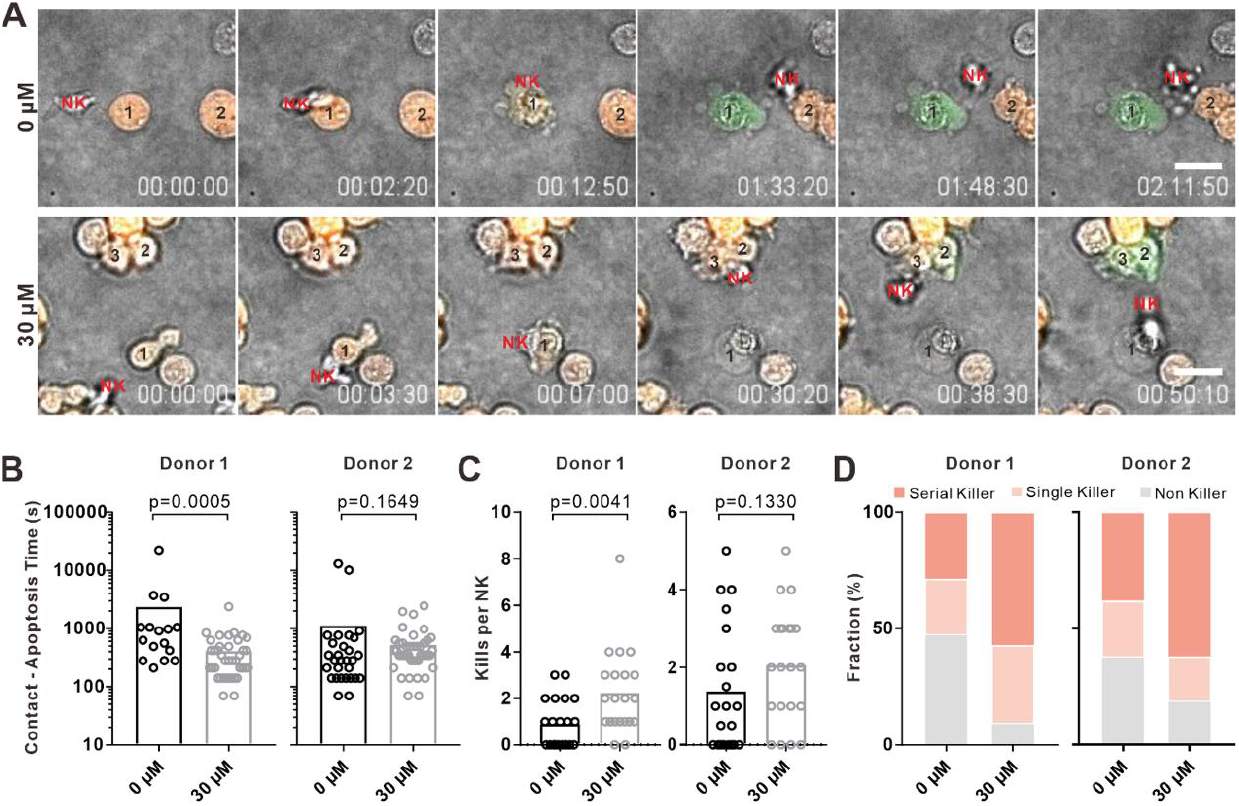
Verbenalin shortened the time required for killing and increases average kills per NK. Primary human NK cells were stimulated with IL-2 in presence of Verbenalin with indicated concentrations for 3 days prior to experiments. Target cells (K562-pCaspar) were embedded in collagen and NK cells were added from top. Killing events were visualized at 37°C every 70 sec. (A) NK cells make multiple contacts with target cells. One representative NK cell from each condition is shown. NK cells (marked in red) were not fluorescently labeled. The target cells in contact with the corresponding NK cell are numbered. Scale bars are 20 μm. (B) Verbenalin shortens the time required for killing. The period from NK/target contact to target apoptosis was quantified. (C) Verbenalin-treatment enhances number of target cells killed per NK cell. (D) Fraction of serial killers is elevated by Verbenalin treatment. Fraction of serial killer (one NK killed more than one target cell), single killer (one NK killed only one target cell) and non-killer (NK cells did not kill any target cells) for each donor was analyzed. Results are from two donors. 21 NK cells were randomly chosen from each condition. Mann-Whitney test was used for statistical analysis.

## 4. Discussion

Uncontrolled immune responses induced by infection is often correlated with life-threatening consequences such as respiratory failure due to lung damage and cytokine-storm syndrome. In this process, exacerbated inflammatory responses initiated by the innate immunity play an essential role. Thus, early interventions minimizing undesired inflammation without sabotaging immune responses to fight pathogens are of great significance to achieve optimal clinical outcome. In this work, we demonstrate that the extract of *V. officinalis*, a medical herb with a long history of utilization in traditional Chinese medicine and alternative medicine in western countries, significantly reduces viral infection-induced acute lung damage as well as release of proinflammatory cytokines. At the same time, administration of low dose of VO-extract can considerably enhance NK activation in response to viral infection. In addition, we have identified that Verbenalin, a biologically active constituent from *V. officinalis*, substantially elevated NK cell-mediated cytotoxicity by shortening the time required for killing and consequently enhancing the frequency of serial killers. These findings suggest *V. officinalis* and Verbenalin as promising therapeutic agents for early intervention to protect lung function, avoid cytokine storm, and facilitate clearance of virus-infected cells.

The use of the *V. officinalis* herb in modern phytotherapy is grounded in its use in folk medicine across different continents including Europe, Asia, America and Australia ^**Error! Reference source not found.**^. Recently, a newly developed formula Xuanfei Baidu composed of thirteen medical herbs including *V. officinalis* has shown very positive clinical outcome treating COVID-19 patients ^**Error! Reference source not found.**^. *V. officinalis* has traditionally been used to treat topical inflammation^[30]^ and chronic generalized gingivitis ^**Error! Reference source not found.**^.

In this study, we demonstrated that VO-extract could relieve lung injury induced by influenza virus A. The effect of VO extract can be a combined result from various levels. It is reported that treatment of *V. officinalis* inhibits the replication of respiratory syncytial virus^**Error! Reference source not found.**^, suggesting that active constituents of *V. officinalis* can inhibit viral replication or induce destruction of virus. We administrated VO-extract intragastrically and the concentration in lung might reach the levels to affect viral replication to some extent. In addition, it is reported that treatment of active constituents of *V. officinalis in vitro* increases phagocytotic activity of neutrophils ^[32]^. We postulate that viral particles in lungs can be removed more efficiently by neutrophils in the VO-extract-treated group.

Massive infiltration of innate immune cells, especially neutrophils and macrophages, into the infected lung tissue is required to remove viral particles, however reactive oxygen species (ROS), nitric oxide (NO), IL-6 and TNF-α released by these cells can lead to damage of endothelial/epithelial cells ^[33]^. Proinflammatory cytokines are the triggers to recruit these cells to inflammation sites. We found that the levels of TNF-α, IL-1β and IL-6 in serum were reduced when treated with VO-extract. TNF-α can be released by M1-type macrophages and T cells ^[35]^, mainly regulated by the NF-κB pathway ^[36]^. IL-1β is released most commonly by monocytes, macrophages, and mast cells; however, non-immune cells, such as neuronal and glial cells, can synthesize and release IL-1β during cell injury or inflammation ^[37]^. Sentinel cells of the innate immune system (macrophages and monocytes) are a major source of IL-1β, but many other cell types, including epithelial cells, endothelial cells, and fibroblasts, can also produce IL-β. IL-1β is regulated by the NF-κB, c-Jun N-terminal kinase (JNK), and p38 MAPK pathways^**Error! Reference source not found.**^. IL-6 can be released by myocardial and immune cell ^[39]^, mainly triggered and regulated by signalling pathways like NF-κB and MAPK ^[40]^. It is reported that total glucosides of *V. officinalis* attenuate chronic nonbacterial prostatitis in rat model by reducing the release of IL-2, IL-1β and TNF-α in prostate^[41]^. Verbenalin, a bioactive constituent of *V. officinalis*, can effectively reduce airway inflammation in asthmatic rats by inhibiting the activity of NF-κB/MAPK signalling pathway ^[42]^. These above-mentioned pathways and molecules can be possible targets for *V. officinalis* to regulate proinflammatory cytokine release.

In this work, we observed that the duration from establishment of NK/target contact to the destruction of target cells was shortened for Verbenalin-treated NK cells relative to their counterpart. To successfully execute their killing function, NK cells need to identify their target cells using surface receptors, followed by formation of a tight junction termed the immunological synapse between the NK cell and the target cells. Consequently, lytic granules (LG) in NK cells, which contain cytotoxic protein such as pore-forming protein perforin and serine proteases granzymes, are enriched and released at the IS to induce destruction of target cell^[43]^. Thus, the duration required for killing is determined by at least five steps: engagement of surface activating/inhibitory receptors, formation of the intimate contact, enrichment of lytic granules, release of lytic granules, and uptake of cytotoxic proteins by target cells.

NK cells express activating receptors and inhibitory receptors to govern NK activation^[45]^. Engagement of activating receptors, such as NKp46, NKp30, NKp44 and NKG2D, triggers signaling pathways for NK activation^[46]^. Inhibitory receptors engage with major histocompatibility complex (MHC) Class I molecules, which are expressed on all healthy self-cells. Loss or down-regulation of MHC-I molecules results in activation of NK cells to initiate the corresponding killing processes ^[47]^. Formation of the intimate contact between NK cells and target cells are largely dependent on LFA-1/ICAM-1 interaction^[48]^. Enrichment of cytotoxic protein-containing LGs at the synapse is regulated by reorganization of cytoskeleton, especially reorientation of MTOC towards the contact site ^[49]^. LG release requires proper granule docking at the plasma membrane and vesicle fusion with the plasma membrane. Transportation and release of LGs are tightly regulated by SNARE and related proteins ^[50]^. Uptake of cytotoxic proteins by target cells requires Ca^2+^ dependent endocytosis^[52]^. Enhancement in any of the above-mentioned steps could accelerate killing processes. For example, sensitize activating receptors and/or up-regulation of effector molecules down-stream of activating receptors, promote LG accumulation at the IS, reduce dwell time for docking, or enhance efficiency of LG release. To identify which step(s) are affected by verbenalin to accelerate killing requires further investigation.

## 5. Conclusions

The present study demonstrated that VO-extract could relieve lung injury induced by influenza virus A and promote the maturation and activation of lung NK cells. Moreover, our *in vitro* study with primary human NK cells from peripheral blood mononuclear cells further suggested Verbenalin significantly reduced contact time required for killing therefore enhancing total killing events per NK cell. These findings established a direct link between Verbenalin, a bioactive constituent of VO-extract, and killing efficiency of NK cells. It indicates that as a compound, Verbenalin has a promising potential for therapeutical applications fighting against cancer and/or infection.

## Declarations

## Abbreviations

IS: immunological synapse;
LG: lytic granules;
MHC: major histocompatibility complex;
NK: Natural killer;
PBMCs: peripheral blood mononuclear cells;
VO-extract: *Verbena officinalis L*. extract.

## Ethics approval and consent to participate

Research carried out for this study with material from healthy donors (leukocyte reduction system chambers from human blood donors) is authorized by the local ethic committee (declaration from 16.4.2015 (84/15, Prof. Dr. Rettig-Stürmer).

## Consent for publication

Not applicable.

## Availability of data and materials

The datasets used and/or analyzed during the current study are available from the corresponding author upon reasonable request.

## Authors’ contributions

RS, HZ, YW and BQ conceived this study. XZ, RZ and AY performed the experiments. RS conducted the network pharmacology analysis. ZX conducted the molecular docking verification. RS and BQ wrote the manuscript. RS, XZ and RZ edited pictures. YW and XD revised the manuscript. All authors read and approved the final manuscript.

## Acknowledgments

We thank the Institute for Clinical Hemostaseology and Transfusion Medicine at Saarland University for providing donor blood; Carmen Hässig, Cora Hoxha, and Gertrud Schäfer for excellent technical help, Markus Hoth for inspiring discussion and continuous support, as well as K562-pCasper cells (with Eva C. Schwarz).

## Supplementary Information

**Supplementary Table 1.**
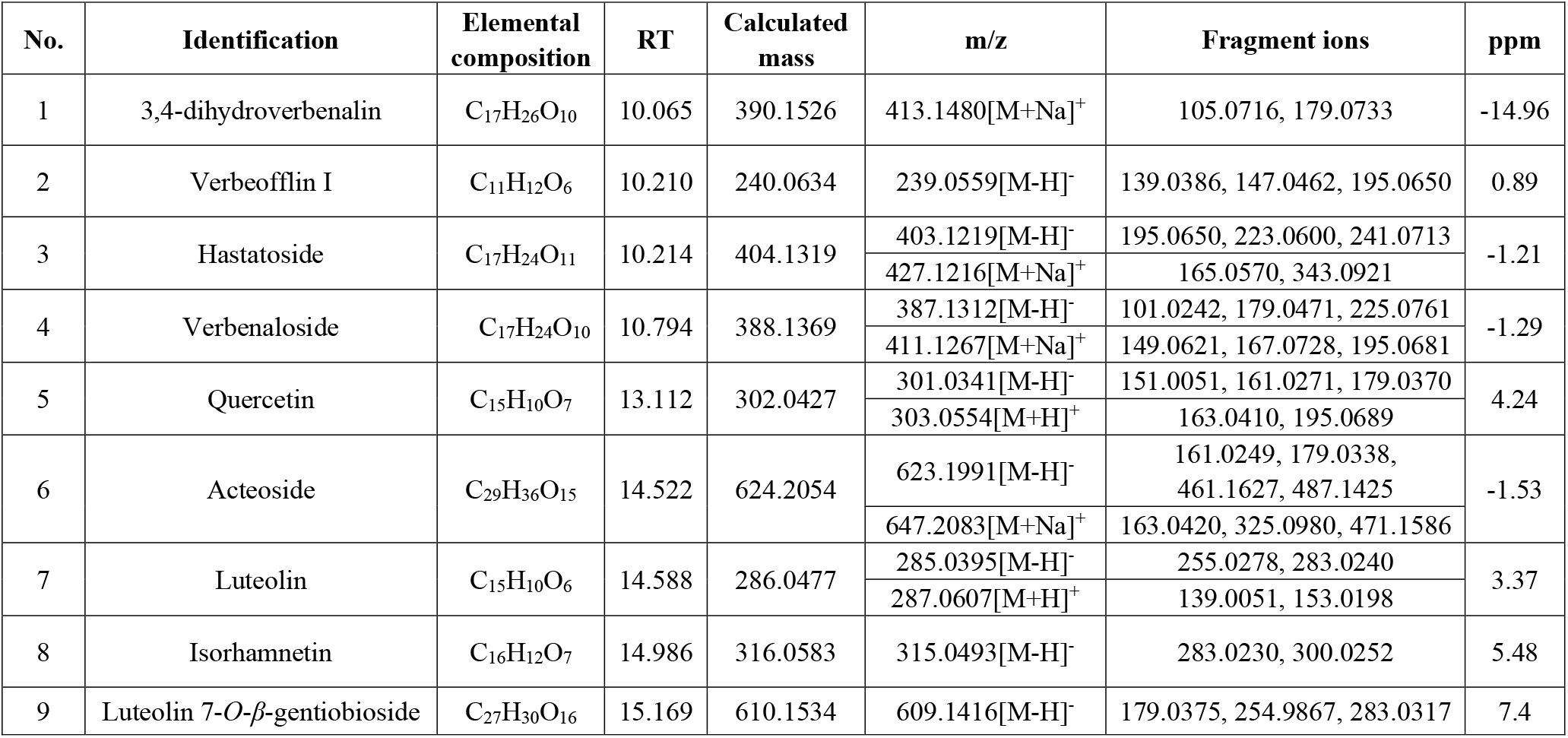
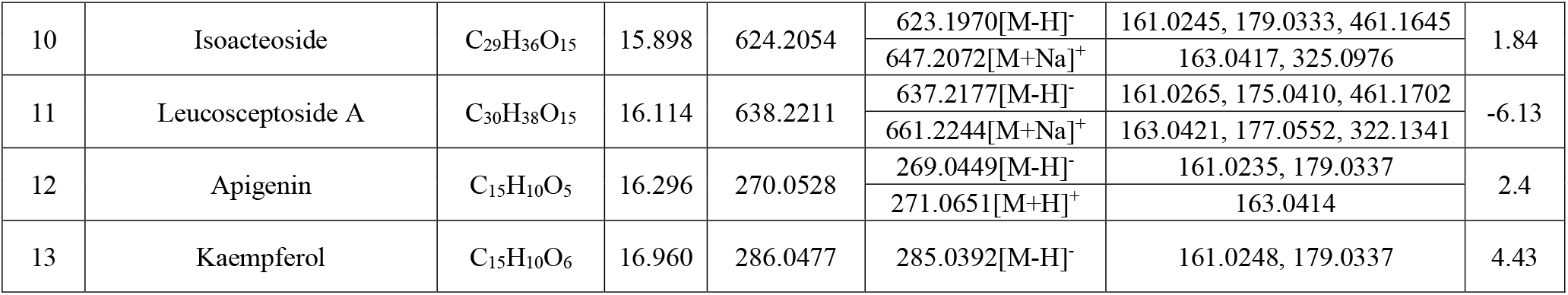
Chemical constituents VO-extract detected by UPLC-Q-TOF-MS.

**Supplementary Movie 1. Verbenalin-treatment reduces time required for killing and enhances average kills per NK.** Primary human NK cells were stimulated with IL-2 in presence of Verbenalin with indicated concentrations for 3 days prior to experiments. Target cells (K562-pCaspar) were embedded in collagen and NK cells were added from top. Killing events were visualized at 37°C every 70 sec. One representative NK cell from each condition (0 μM vs 30 μM) is shown. NK cells were not fluorescently labeled and marked with blue tracks. The target cells in contact with the corresponding NK cell are numbered.

